# A Single-Cell Immune Atlas of Triple Negative Breast Cancer Reveals Novel Immune Cell Subsets

**DOI:** 10.1101/566968

**Authors:** Si Qiu, Ruoxi Hong, Zhenkun Zhuang, Yuan Li, Linnan Zhu, Xinxin Lin, Qiufan Zheng, Shang Liu, Kai Zhang, Mengxian Huang, Kaping Lee, Qianyi Lu, Wen Xia, Fei Xu, Xi Wang, Jun Tang, Xiangsheng Xiao, Weidong Wei, Zhongyu Yuan, Yanxia Shi, Yong Hou, Xiuqing Zhang, Jian Wang, Huanming Yang, Qimin Zhan, Bo Li, Shusen Wang

**Affiliations:** Sun Yat-sen University Cancer Center, State Key Laboratory of Oncology in South China, Collaborative Innovation Center of Cancer Medicine, Guangzhou 510060, China; BGI-Shenzhen, Shenzhen 518103, China; BGI-GenoImmune, BGI-Shenzhen, Wuhan 4300794, China; China National GeneBank, BGI-Shenzhen, Shenzhen 518120, China; School of Biology and Biological Engineering, South China University of Technology, Guangzhou 510006, China; BGI Education Center, University of Chinese Academy of Sciences, Shenzhen 518083, China; Guangdong Enterprise Key Laboratory of Human Disease Genomics, Shenzhen 518103, China; Shenzhen Key Laboratory of genomics, Shenzhen 518103, China; James D. Watson Institute of Genome Sciences, Hangzhou 310008, China; Senior author, Key laboratory of Carcinogenesis and Translational Research (Ministry of Education/Beijing), Laboratory of Molecular Oncology, Peking University Cancer Hospital & Institute, Beijing 100142, China

**Author notes:** These authors contributed equally to this work (Si Qiu, Ruoxi Hong, Zhenkun Zhuang, Yuan Li, Linnan Zhu). Correspondence: Shusen Wang, Bo Li, Qimin Zhan.

## Abstract

Triple-negative breast cancer (TNBC) represents the most aggressive breast cancer subtype, which recently attracts great interest for immune therapeutic development. In this context, in-depth understanding of TNBC immune landscape is highly demanded. Here we report single-cell RNA sequencing results of 9683 tumor-infiltrated immune cells isolated from 14 treatment naïve TNBC tumors, where 22 immune cell subsets, including T cells, macrophages, B cells, and DCs have been characterized. We identify a new T cell subset, CD8^+^CXCL8^+^ naïve T cell, which associates with poor survival. A novel immune cell subset comprised of TCR^+^ macrophages, is found to be widely distributed in TNBC tumors. Further analyses reveal an up-regulation of molecules associated with TCR signaling and cytotoxicity in these immune cells, indicating TCR signaling activation. Altogether, our study provides a valuable resource to understand the immune ecosystem of TNBC. The novel immune cell subsets reported herein might be functionally important in cancer immunity.

**SIGNIFICANCE:** This work demonstrates a single-cell transcriptome atlas of immune cells in treatment naïve TNBC tumors, revealing novel immune cell subsets. This study provides a valuable resource to understand the immune ecosystem of TNBC, which will be helpful for the immunotherapeutic strategy design of TNBC.

## INTRODUCTION

Cancer immunotherapies are revolutionizing cancer treatment landscape. Current checkpoint blockade therapies mainly function to rescue T cells from exhaustion or eliminate T regulatory cells (Treg). Emerging evidences indicate that myeloid cells, such as macrophages, are highly influential on other cell populations in the tumor microenvironment (TME), including cancer cells as well as immune cells (1–3). Whereas most myeloid cells promote cancer outgrowth, others display potent anti-tumour activity (3). Targeting the myeloid cell population has attracted increasing interest in recent years (4, 5). While checkpoint blockade therapies have demonstrated remarkable clinical activity in many cancer types, the biological determinants of response to these agents remain incompletely understood. Several factors such as tumor neoantigens, mutation burden, tumor-infiltrating lymphocytes (TILs) levels, and PD-L1 expression have shown correlation with response, but other factors from the microenvironment also have profound influences (6). The design of novel cancer immunotherapy strategies and the identification of effective clinical biomarkers require deep understanding of this ecosystem.

T cells are the most abundant and best-characterized population in the TME of solid tumors (7, 8). CD4^+^ helper T cells and CD8^+^ cytotoxic T cells can exert anti-tumor effect by targeting antigenic tumor cells, and levels of activated CD8^+^ T cells are predictive of good prognosis in several cancers (9, 10). However, the tumor microenvironment can develop various mechanisms to suppress T cell responses and facilitate cancer cell survival. These mechanisms may involve Tregs, which secrete immunosuppressive cytokines, and myeloid and stromal cells, which modulate immune check points by activation of co-inhibitory receptors (e.g., PD-1, Tim-3, and CTLA-4) on T cells, driving T cell dysfunction and exhaustion (11, 12). Understanding the mechanisms of TME induced T-cell dysfunction should assist on the development of promising combinatorial immunotherapies to improve the clinical efficacy of current immune checkpoint blockades.

Tumor-associated macrophages (TAMs) are another key component of the TME with a dual supportive and inhibitory influence on cancer growth (13, 14). Customized functional model divides macrophages into to two categories: classical M1 and alternative M2 macrophages. The M1 macrophage is involved in the inflammatory response, pathogen clearance, and antitumor immunity. In contrast, the M2 macrophage influences an anti-inflammatory response, wound healing, and pro-tumorigenic properties (15). TAM phenotypes are highly plastic, and recent reports show that the model distinguishing between classically polarized anti-tumor M1 and alternatively polarized pro-tumor M2 subtypes incompletely accounts for the phenotypic diversity in vivo (16). Azizi *et al* found that M1 and M2 gene signatures are positively correlated in the myeloid populations in breast cancer tumors (17).

Triple-negative breast cancer (TNBC) represents up to 20% of all breast cancers. TNBC tumors are typically more aggressive and difficult to treat than hormone receptor-positive tumors, and are associated with a higher risk of early relapse. The lack of estrogen receptor, progesterone receptor, and HER2 expression precludes the use of targeted therapies, and the only approved systemic treatment option is chemotherapy. Responses to chemotherapy occur, but are often short lived and frequently accompanied by considerable toxicity. Gene profiling studies reveal that TNBCs are highly heterogeneous and a large proportion of them demonstrate DNA Repair Deficiency Signature (18–20). Recent data showed impressive activity of PD-1/PD-L1 blockade therapy in metastatic TNBC patients who were chemotherapy naïve, suggesting early intervention of immunotherapy can bring more benefit (21). Clinical trials applying checkpoint inhibitors in the neo-adjuvant setting of TNBC are ongoing. Although need to be confirmed in a larger cohort, these results are consistent with the notion that immunotherapy agents are most efficient at low tumor burden and in patients naïve of immune-modulatory chemotherapy agents. To better understand the immune ecosystem of TNBC, we analyzed the full-length single-cell RNA sequencing data of 9,683 tumor-infiltrated immune cells isolated from treatment naïve TNBC tumors. We identified 22 unique immune cell subsets, including T cells, macrophages, B cells, and DCs. Using combined expression and TCR-based analyses, we were able to indicate the function and developmental path of T cell subsets. We found a novel T cell subset, CD8^+^CXCL8^+^ naïve T cell, and revealed its possible tumor-promoting function. Three subpopulations of tumor infiltrated CD3^+^/CD4^−^/CD8^−^ double-negative (DN) T cells were identified. We also demonstrated the co-expression pattern of M1/M2 signature on single-cell macrophages and demonstrated the widely existence of TCR^+^ macrophages. This is the first time that TCR^+^ macrophages were studied on single-cell transcriptome level.

## RESULTS

### Tumor Characteristics and Single-cell Transcriptome Profiling of Immune Cells

To uncover the complexity of immune ecosystem in TNBC, we performed deep single-cell RNA sequencing on immune cells flow-sorted from the primary tumors of 14 treatment naïve TNBC patients based on the expression of CD45 (Fig. 1a and Supplementary Fig. 1a). After filtering out cells with low quality, we obtained 9,683 CD45^+^ cells with an average of 14.80 million uniquely mapped reads per cell (Supplementary Fig. 1b and Supplementary Table 1). This sequencing depth assured the detection for genes with low expression level, allowing reliably profiling of cytokines and transcription factors in immune cells (Supplementary Fig. 1c).

**Figure 1.**
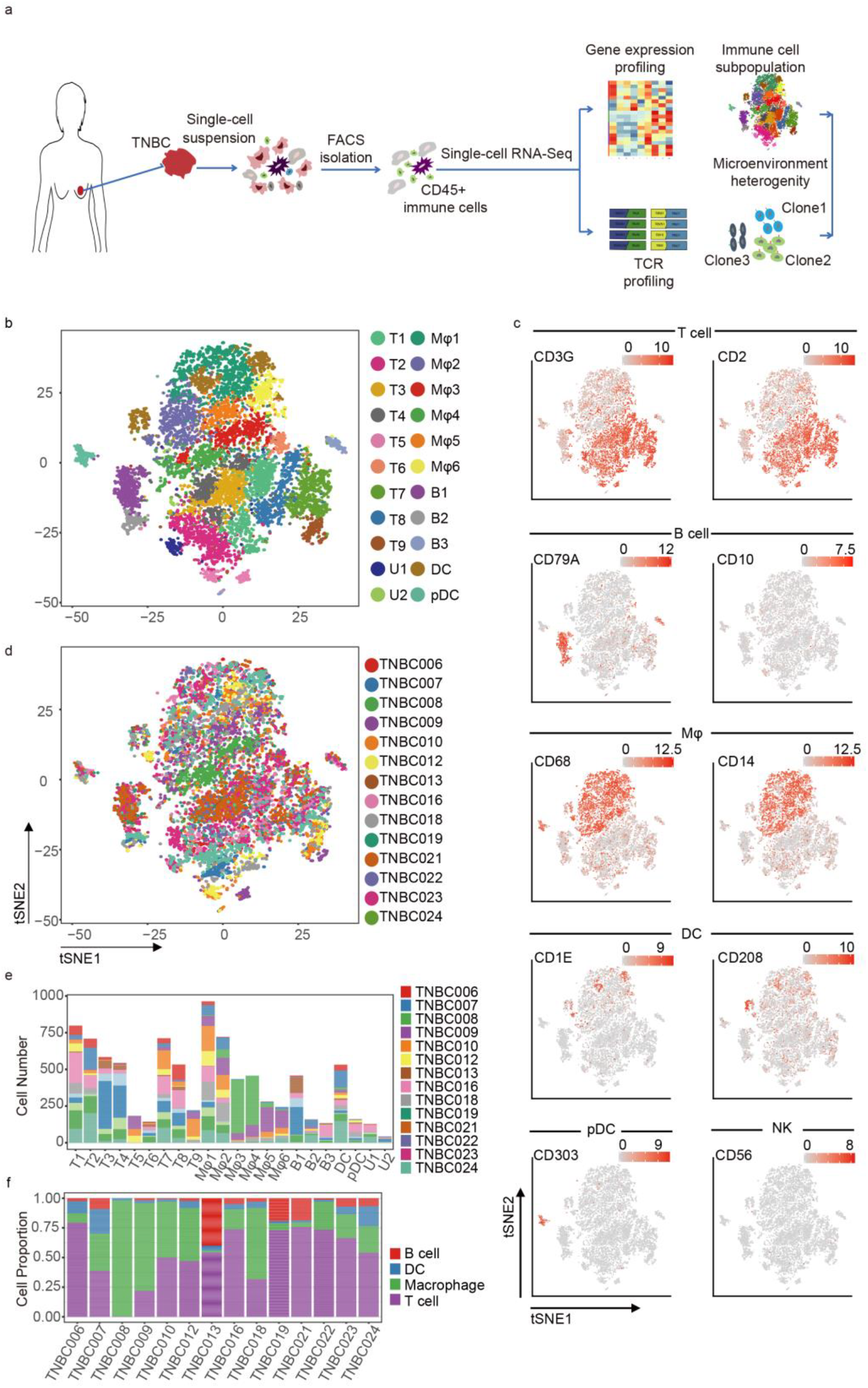
Program Design and Immune Landscape of TNBC. **a**, Flow chart of the experimental design. Single-cell RNA sequencing was applied to CD45^+^ cells derived from tumor tissue. The scRNA-seq data were then subjected to gene expression profiling and TCR profiling. **b**, Two-dimensional t-SNE plot of 9,683 CD45^+^ single cells from 14 TNBC patients. Each point represents one single cell, colored according to cell cluster. **c**, Cells colored by normalized expression (log_2_(TPM+1)) of immune cell type classification markers on the t-SNE map. **d**, Similar to **b**, colored according to patient. **e**, Bar plot shows the fractions of cells from different patients in each cell cluster, colored by patient. **f**, Bar plot shows the cell type fractions in each patient’s tumor-infiltrating immune cells, colored by cell type. T1-T9, T cell clusters. MΦ1-MΦ6, macrophage clusters. B1-B3, B cell clusters. DC and pDC, DC clusters. U1 and U2, undefined clusters.

### The Immune Landscape of TNBC

To reveal the immune cell populations in TNBC, we performed unsupervised clustering using Seruat method on CD45^+^ cells and obtained 22 cell clusters, which were then visualized by t-SNE algorithm (22) (Fig. 1b). We identified the immune cells as T cells, B cells, macrophages and DCs based on the expression of classic cell type markers as well as reference component analysis (23) (Fig. 1c and Supplementary Fig. 1d). With this approach, nine clusters were annotated as T cell clusters, six as macrophage clusters, three as B cell clusters, two as DC clusters, and also there remained two clusters that could not be well determined (Fig. 1b and Supplementary Fig. 1e). T cells were the predominant immune cell population, accounting for 46.58% of the total amount, followed by macrophages (35.71%), B cells (8.23 %) and DCs (7.71%). The existence of CD8^+^ T cells, CD4^+^ regulatory T cells, B cells and macrophages were confirmed by immunohistochemical staining (Supplementary Fig. 1f). The two DC clusters could be further categorized as one classical DC subset that expressed *CD1c* and *Dectin 1* (*CLEC7A*), and one plasmacytoid DC subset specifically expressing *IL3RA* and *CD303* (*CLEC4C*)(24). For the B cell clusters, all three clusters are CD10^−^ mature B cells(25). Most of the clusters were consisted of cells from multiple patients, indicative of common immune traits among patients (Fig. 1d). Besides, the proportion of different immune cell types and clusters varied across patients, consistent with descriptions in previous researches (Fig. 1e,f) (17).

### Subtype Analysis of Tumor Infiltrated T cells

T lymphocytes are the most abundant and well-studied immune cell population in the microenvironment of solid tumors. We sought to characterize infiltrated T cells in TNBC tumors. Unsupervised clustering of CD45^+^ cells identified nine T cell clusters, including three clusters of CD4^+^ T cells, and six clusters of CD8^+^ T cells (Fig. 2a). Distinct signatures of T cell clusters were revealed by expression patterns of known T cell functional markers (cytotoxic, naïve, regulatory or exhausted) and differentially expressed genes (DEGs) among clusters (Fig. 2b, c and Supplementary Table 2). Among the CD8^+^ clusters, T2 and T4 were suggested as naïve T cell clusters for their specific expression of *CCR7, SELL*, and *LEF1* (26). T1 and T6 showed high expression of exhaustion markers *HAVCR2* (*TIM-3*)*, TIGIT*, and *LAG3*(27), indicative of exhausted CD8^+^ T cell clusters. T3 and T5 were characterized by high expression of genes associated with cytotoxicity, including *IFNG*, *PRF1*, *GZMA* and *GZMB*, and meanwhile expressed low levels of exhaustion markers, representing cytotoxic T cell clusters (Fig. 2b).

**Figure 2.**
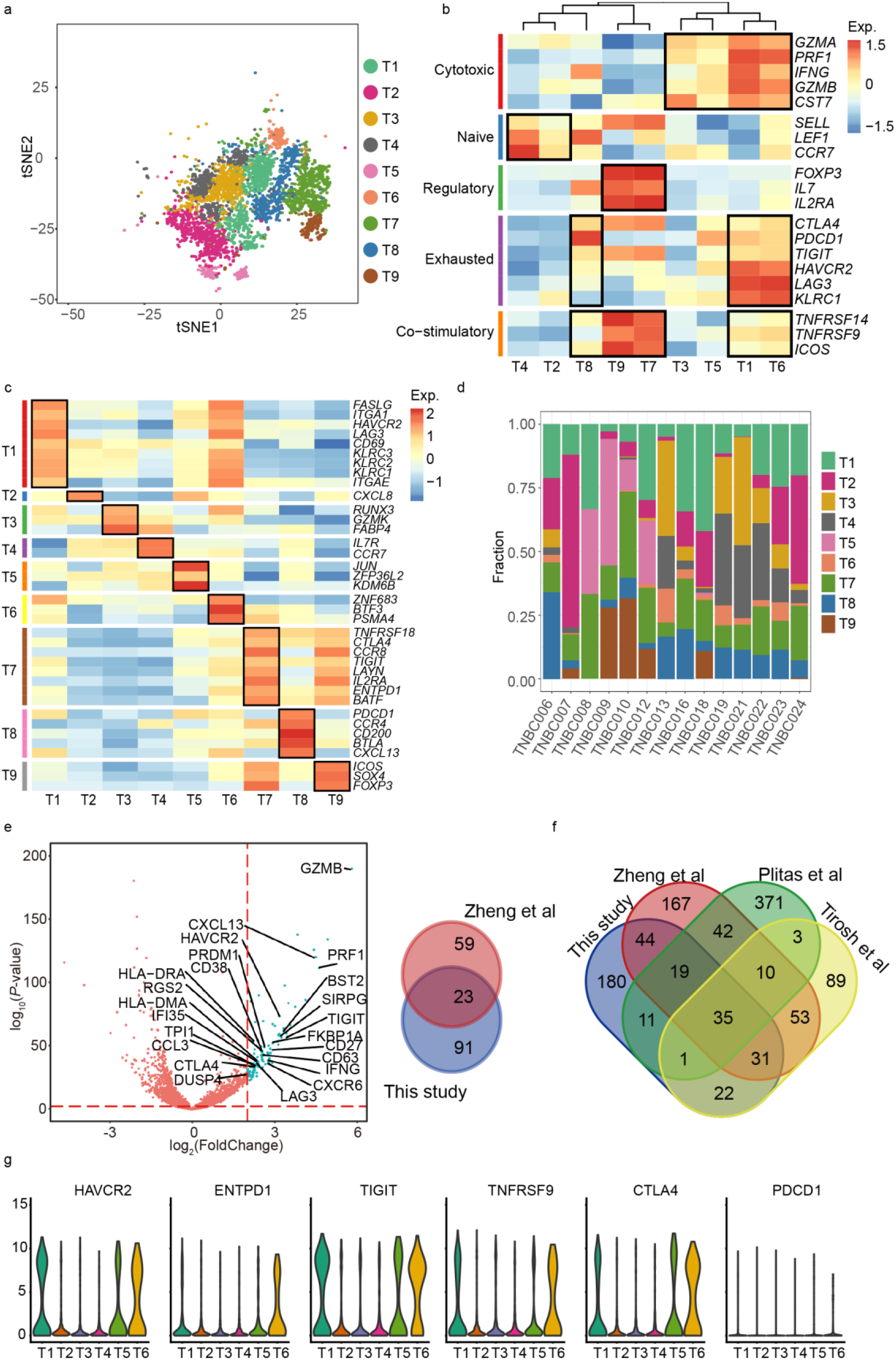
Clustering and Functional Analysis of T Cells. **a**, Two-dimensional t-SNE plot of 9 identified T cell clusters. Each point represents one single cell, colored according to cell cluster. **b**, Z-score normalized mean expression (log_2_(TPM+1)) of selected T cell function-associated genes in each T cell cluster. Black boxes highlight the patterns defining known T cell subtypes. **c**, Z-score normalized mean expression (log_2_(TPM+1)) of most representative markers of each T cell cluster. Black boxes highlight the markers for specific cluster. **d**, The fractions of nine T cell clusters within each patient. **e**, The volcano plot in the left panel shows differentially expressed genes in tumor-infiltrating exhausted T cells. Each blue dot represents a gene with P-value < 0.01 and log_2_FC ≥ 2. Totally 23 genes overlap with previous HCC study by Zheng et al. (2017), which are marked with gene names. The right panel shows the overlap of exhausted T cell signature genes identified in this study with those from previous study by Zheng et al. (2017) (p < 2.2e-64), determined by hypergeometric test. **f**, Venn graph showing the overlap of Treg signature genes identified in this study with those from previous studies by Zheng et al. (2017), Plitas et al. (2016), and Tirosh et al. (2016). **g**, The expression levels of exhausted markers in CD8^+^ T cell clusters.

For the three CD4^+^ clusters, T7 and T9 exhibited remarkable regulatory T cell (Treg) features for high expression of *FOXP3* and *IL2RA* (*CD25*). The remaining CD4^+^ cluster T8 was marked by high levels of *CXCL13*, *CD200*, *BTLA*, *PDCD1* (*PD-1*), and *CTLA4*, consistent with previously reported dysfunctional T helper cell (Th) features (28, 29). Both CD8^+^ and CD4^+^ T cell clusters exhibited distinct distributions among patients (Fig. 2d).

### Genes Uniquely Expressed by Exhausted CD8^+^ T Cells and Tumor-infiltrated Tregs in TNBC

Current immune checkpoint blockade therapies mainly target on exhausted CTL or Tregs. To uncover potential therapeutic targets for immunotherapy of TNBC, we sought to investigate genes uniquely expressed by these two immunosuppressive T cell subsets. By comparing the expression profiles of exhausted CD8^+^ clusters (T1 and T6) with non-exhausted clusters (T2, T3, T4 and T5) using R package limma, we obtained a set of 114 exhaustion-specific genes (adjusted P < 0.01 and log_2_FC ≥ 1, Supplementary Table 3), including multiple known exhaustion markers, such as *TIGIT*, *HAVCR2*, *LAG3*, *CTLA4*, and *KLRC1*(12) (Fig. 2e). Among these genes, 23 were also demonstrated in the exhausted CD8^+^ T cell specific gene list by a previous study of liver cancer (30) (Fig. 2e; Supplementary Table 4). Notably, tumor-infiltrating CD8^+^ T cells in our TNBC tumors expressed low level of PDCD1 (PD-1), while high levels of other exhaustion markers such as HAVCR2, TIGIT, and CTLA4 were detected (Fig 2g). While both exhausted T cell clusters showed high levels of exhaustion marker expression, T1 was also characterized by tissue-resident memory T (TRM) cell feature for high expression of TRM marker genes, such as *CD69*, *ITGAE* (*CD103*) and *ITGA1 (31)*.

The same method was applied to tumor-infiltrating Tregs and a total of 343 genes uniquely expressed by Tregs were identified (Supplementary Table 5). The TNBC Treg-specific genes largely overlapped with those identified in previous studies in liver cancer, melanoma and breast cancer (30, 32) (Fig. 2f).

### Pseudotime State Transition of T cells and a “Pre-exhaustion” T Cell Subset

Pseudotime analysis provides us a method to depict the T cell developmental trajectories that correspond to biological processes such as activation or exhaustion, as recently suggested (30). To probe into the T cell functional state transition in TNBC, we used the Monocle2 algorithm to order CD8^+^ and CD4^+^ T cells in pseudotime based on transcriptional similarities (33). The CD8^+^ T cell trajectory began with cells of naïve clusters T2 and T4, which were separately located at two different branches, followed by cytotoxic clusters T3 and T5, and terminated with exhausted clusters T1 and T6 (Fig. 3a). The enrichment of exhausted T cells at the late period of the pseudotime was in line with previous analysis (30, 31), indicating the CD8^+^ T cell developmental process through activation to exhaustion. Similarly, we analyzed the CD4^+^ T cells’ differentiation trajectory, in which Treg clusters T7 and T9 were located at the other end of the exhausted CD4^+^ Th cluster T8, demonstrating distinct functional states among these T cell subsets (Fig. 3b).

**Figure 3.**
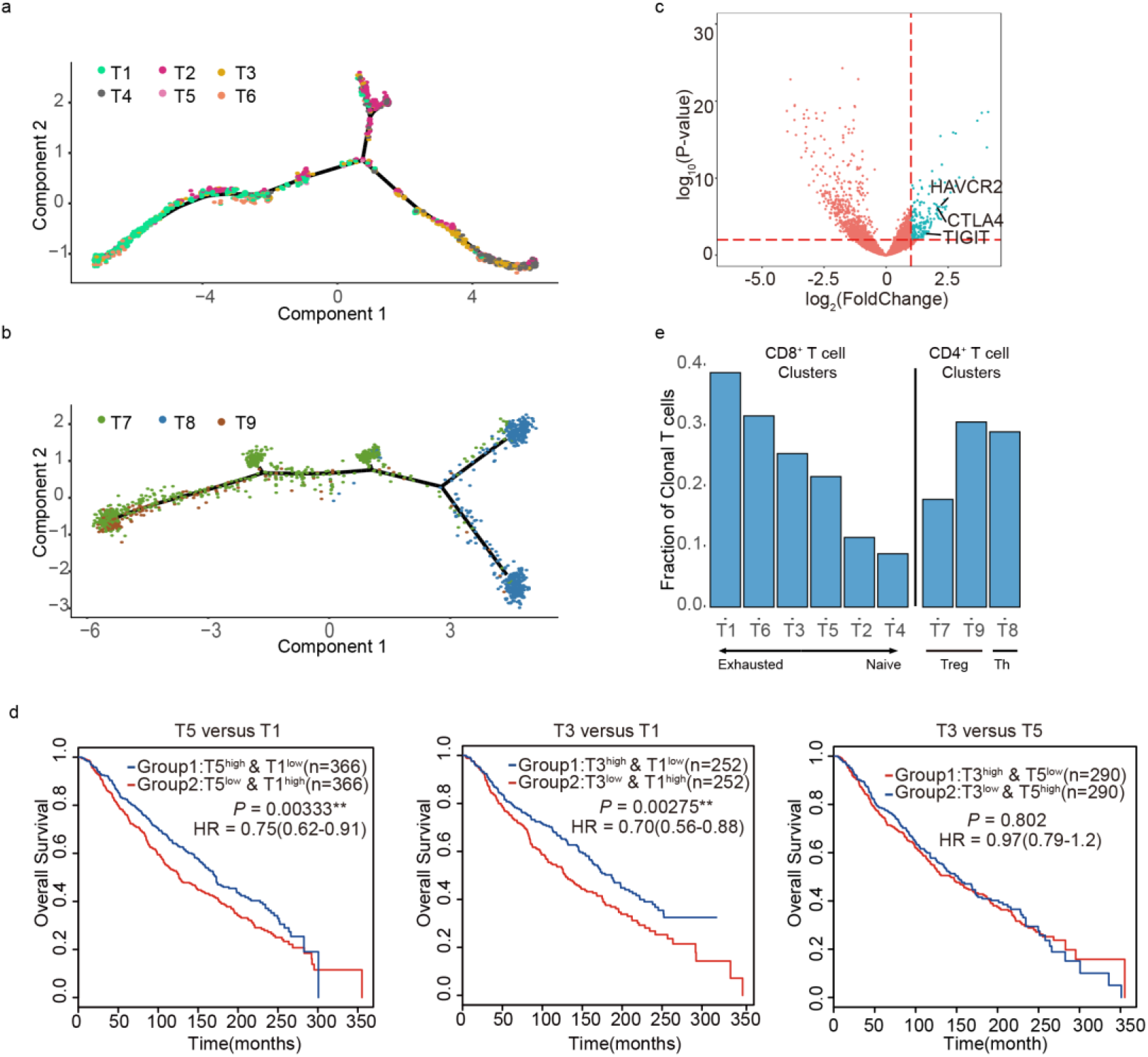
T Cell Trajectory Analysis Reveals a “Pre-exhaustion” T Cell Subset. **a**, The trajectory of CD8^+^ T cells shows CD8^+^ T cell state transition in a two-dimensional space defined by Monocle2. Each point represents one single cell, colored according to cell cluster. **b**, Similar plot as **a** for CD4^+^ T cell clusters. **c**, Differential genes between T3 and T5. Each blue dot represents a gene with P-value < 0.01 and log2FC ≥ 2. **d**, The Kaplan–Meier overall survival curves of METABRIC patients grouped by the gene signature score (see Method) of T5 and T1(left panel), T3 and T1 (middle panel), T3 and T5 (right panel). The high and low groups are divided by the median value of gene signature score. *P*-value was calculated by multivariate Cox regression. HR, hazard ratio. **e**, Comparison between proportions of clonal cells in each T cell cluster.

As shown by the trajectory analysis of CD8^+^ T cells, T5 appeared to be an intermediate functional state, locating between cytotoxic cluster (T3) and exhausted clusters (T1 and T6). Comparison of the expression patterns of T5 and T3 revealed a set of 259 genes with elevated expression in T5, including exhausted marker genes *CTLA4*, *HAVCR2*, and *TIGIT* (Fig. 3c). Whereas survival analysis of the BC cohort from METABRIC revealed that patients expressing T5^high^ T1^low^ or T3^high^ T1^low^ signature were associated with significantly better survival than patients expressing T1^high^T5^low^ or T1^high^T3^low^ signature, respectively. Also, patients expressing high T3 and low T5 signature showed comparable survival when compared with high T5 and low T3 signature patients (Fig. 3d). This survival disparity suggested that although T5 showed an elevated expression of exhausted markers, its signature was associated with more favorable prognosis than the exhausted T1 signature, consistent with the description of a “pre-exhaustion” T cell state that have been suggested in lung cancer and liver cancer (30, 31). Genes involved in the cellular senescence pathway, such as *KDM6B* and *ZFP36L2* (Supplementary Table 2), were highly expressed by cells of T5, indicating a possible mechanism that triggers T cell exhaustion (34).

### Clonal Enrichment of Exhausted T Cells and Tregs in TNBC Microenvironment

TCR analysis provides another approach to gain insight into the various states of T cells. The diversity of TCR sequences is pivotal for recognizing viral antigens or tumor-specific neoantigens presented by the major histocompatibility complex (MHC) on antigen-presenting cells (APCs). While the TCR repertoire is enormous due to the large amount of TCRs and random recombination, identical TCR sequences can indicate T cell clonal expansion. Our single-cell RNA-seq data allowed us to track the lineage of each T cell based on their full-length TCR α and β sequences assembled by the TraCeR method (35). Full-length TCRs with both α and β chains were obtained for 3,315 T cells from 14 patients, among which 2,311 harbored unique TCRs and 1,004 harbored repeatedly used TCRs, implying clonal expansion. Patterns of clonal expansion were detected at varying degrees in different T cell clusters. While only approximately 10% CD8^+^ cells of naïve clusters (T2 and T4) harbored clonal TCRs (those whose α and β TCR pairs were shared by at least two cells), the percentage reached over 30% in exhausted CD8^+^ T cell clusters (T1 and T6). For the CD4^+^ clusters of Treg and exhausted Th, 17.82% to 30.61% of clonal CD4^+^ T cells were observed (Fig. 3e). Thus, identifying clonal TCRs at single-cell level verified the previously suggested activation and exhaustion status of different T cell clusters in TNBC microenvironment.

### A CXCL8 Producing CD8^+^ Naïve T Cell Subset Associated with Survival

CXCL8 production is mostly ascribed to myeloid and epithelial cells, and previous studies have shown that human umbilical cord blood CD4^+^ naive T cells can express CXCL8, but CXCL8-producing T cells become rare in adults (36). Here we observed T2, a new subset of CD8^+^CXCL8^+^ naïve T cells, which was found in 10 of 14 TNBC tumors and accounted for 16.00% of the T cell population (Figure 4a). The high fraction of CXCL8^+^ T cells highlighted the possibility that this T cell subset might have active and potentially significant functions in TNBC microenvironment. By analyzing the T cell function-associated gene expression in T2, we found this subset of cells expressed high levels of naïve markers CCR7 and SELL, and activation marker CD69, but low levels of differentiation markers, including CD57 and KLRG1 (Supplementary Fig. 2a). These CXCL8^+^ T cells expressed low levels of CXCR1 or CXCR2, the receptor for CXCL8, but elevated levels of other chemokines including CCL3, CCL4 and CCL2, when compared with the other naïve T cell cluster T4 (Supplementary Table 6). These chemokines might work corporately with CXCL8 in TNBC microenvironment. Interestingly, a recent study revealed that CD4^+^CXCL8^+^ naïve T cells might support tumor growth and lymphoid metastasis via CXCL8 mediated neutrophil migration (37). The tumor promoting effect of the newly identified CD8^+^CXCL8^+^ naïve T cell was validated by survival analysis of an independent METABRIC cohort of breast cancer. This analysis showed that patients with high expression of T2 signature genes and low T4 signature had significantly worse survival compared to those with high T4 and low T2 signature gene expression (*P* = 0.0263, Fig. 4b). This suggests that CXCL8^+^ naïve T cells may associate with poor prognosis in breast cancer, illuminating a potential therapeutic target in TNBC. To investigate the underlying mechanisms of CD8^+^CXCL8^+^ naïve T cells driving tumor aggression, we performed DEG analysis between tumors showing high and low T2 signature using data from the METABRIC cohort. This analysis revealed 75 genes with elevated expression (Log_2_FC≥1) in the T2^high^ group (Supplementary Fig. 2b; Supplementary Table 7). GO biological process enrichment analysis showed these highly expressed genes were generally enriched in the leukocytes (including granulocyte and myeloid leukocyte) chemotaxis and migration pathways (Fig. 4c; Supplementary Table 7). Moreover, this analysis also revealed that MAPK and ERK1/ERK2 cascades were activated in the T2^high^ group (Supplementary Table 7). These results suggested that CD8^+^CXCL8^+^ naïve T cells, a subset of chemokine producing naïve T cells, might promote cancer progression through mediating leukocytes migration to tumor site and activating the MAPK/ERK pathways.

**Figure 4.**
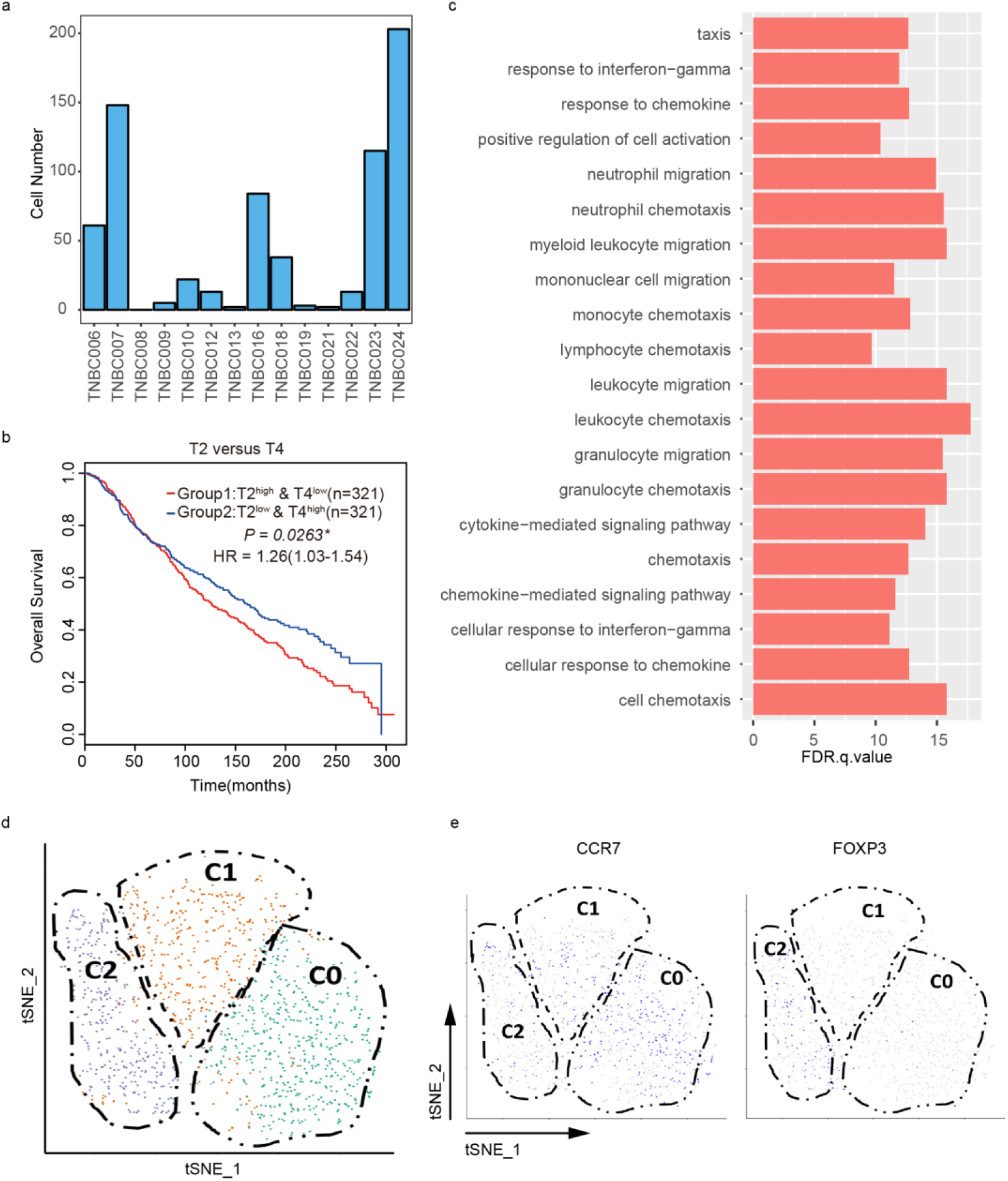
Analysis of CD8^+^ CXCL8^+^ Naïve T cell subset and DNT cells. **a**, The distribution of CD8^+^ CXCL8^+^ Naïve T cell subset (T2) among patients. **b**, The Kaplan-Meier overall survival curves of METABRIC patients grouped by the gene signature score (see Method) of T2 and T4. The high and low groups are divided by the median value of gene signature score. *P*-value was calculated by multivariate Cox regression. HR, hazard ratio. **c**, Results of GO analysis using genes up-regulated in patients with T2^high^ signature. **d**, Two-dimensional t-SNE plot of 3 identified DN cell clusters. Each point represents one single cell, colored according to cell cluster. **e**, Cells colored by normalized expression (log_2_(TPM+1)) of CCR7 and FOXP3 on the t-SNE map.

### Characterizing and Clustering of DN T cells in TNBC Microenvironment

CD3+/CD4-/CD8-double negative (DN) T cells, comprising 1% to 5% of the total T cell population in healthy humans, have been shown to play roles in inflammation and autoimmunity (38, 39). Whereas DN T cells have not been well-characterized in the tumor microenvironment. In this study, we observed substantial amount of tumor-infiltrated DN T cells, taking up 31.0% (1324/4285) of the total amount of T cells (Supplementary Fig. 2c). Unsupervised clustering of these DN T cells revealed three clusters, as visualized in t-SNE map (Figure 4d). C1 expressed high levels of cytotoxic markers like GZMA, GZMB and IFNG (IFN-γ), indicative of an effector-like group. C2 appeared to be a regulatory population expressing high level of FOXP3, IL2RA and CTLA4. DN Tregs have been shown to be associated with autoimmune disease (40). C0 represented a naïve-like population with high CCR7 expression (Figure 4e). TCR-based analyses revealed that 68.5% DN cells harbored at least one paired productive α-β chains while 2.4% harbored paired productive γ-δ chains. In total, 71 TCR pairs were shared with CD4^+^ T cells, 31 with CD8^+^ T cells and 52 within DN T cells. These findings indicate that the tumor-infiltrated DN T cell population may exert various functions and demonstrate clonal accumulation that similar to CD4 or CD8 single positive conventional T cells. The high proportion and diverse subtyping of DN T cells suggests that this somewhat neglected class of T cells may be functionally important in TNBC immunity.

### Characterization of TAM subpopulations in TNBC

For the categorization of the macrophages in TNBC, we first applied the classic macrophage polarization model. Based on the activation state, macrophages can be recognized as anti-tumoral classically activated (M1) cells and pro-tumoral alternatively activated (M2) cells (41). We observed highly positive correlation of M1 and M2 signature gene expressions in all the macrophages we identified (Pearson correlation test, R = 0.624, *P* < 2.20 × 10^−16^, Fig. 5a). Although this M1/M2 co-expressed state in single cell level raised challenge to the classic model of macrophage polarization showing exclusive discrete M1 and M2 activation state, along with recent reports (42, 43), we were able to classify the macrophages into six clusters using Seruat based on their transcriptome features. Totally, we identified six macrophage clusters, each one of which had its uniquely expressed genes (Fig. 5b, Supplementary Table 8).

**Figure 5.**
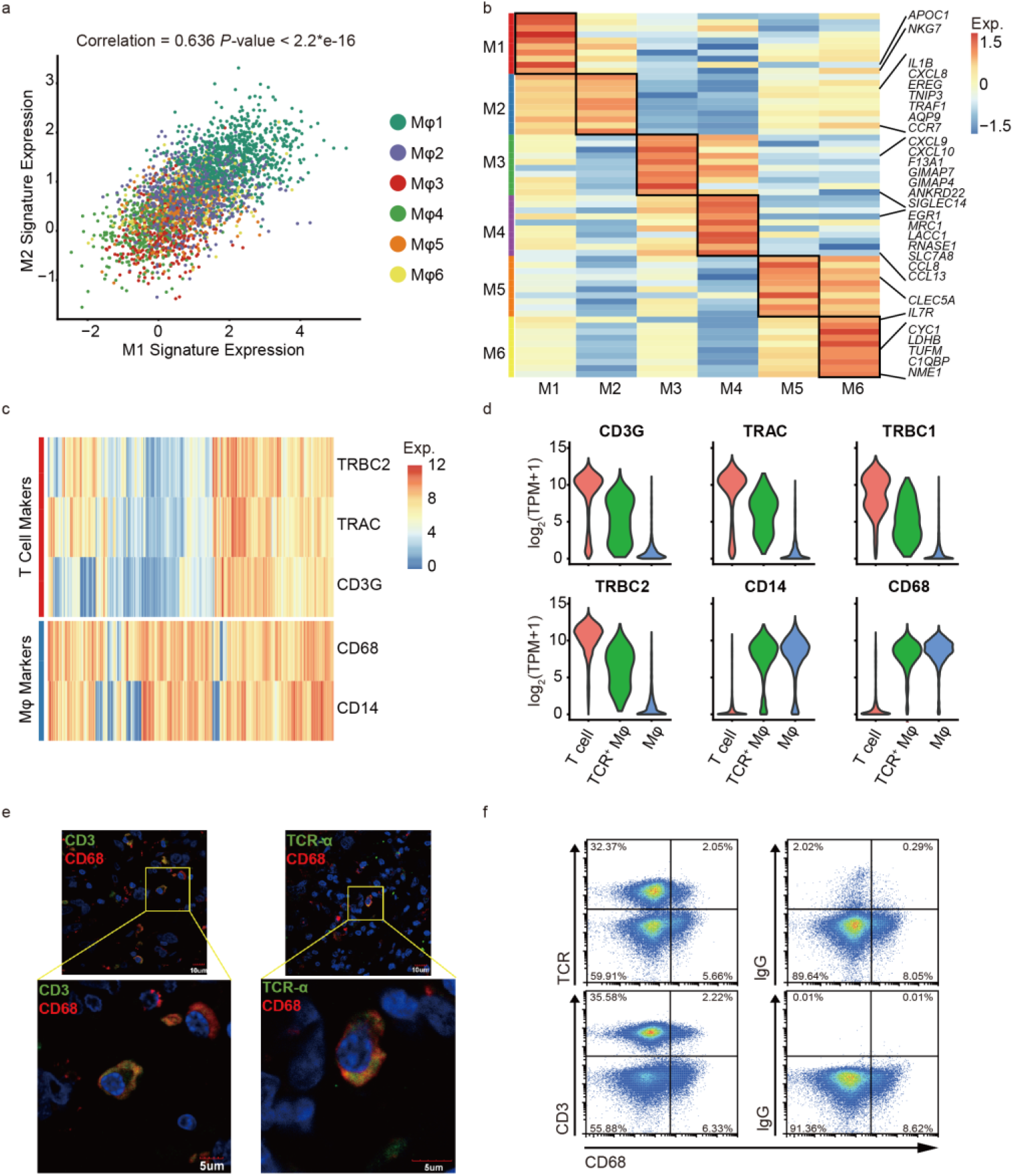
Characteristics of macrophage clusters and TCR^+^ macrophages. **a**, Scatterplot of mean normalized expression (see Method) of M1 and M2 signatures per cell. Each point represents one single cell, colored by clusters. **b**, Z-score normalized mean expression (log_2_(TPM+1)) of representative markers of each macrophage cluster. Black boxes highlight the markers for specific cluster. **c**, The expression (log_2_(TPM+1)) of T cell marker (CD3G), TCR genes (TRBC2 and TRAC) and macrophage markers (CD68 and CD14) in TCR^+^ macrophages (n=382). **d**, Violin plots show the expression (log_2_(TPM+1)) of above genes across T cells, TCR^+^ macrophages and normal (TCR^−^) macrophages. **e**, Immunofluorescence proposes the co-expression of TCR/CD3 and CD68 in paraffin section of breast cancer samples. **f**, Co-expression of TCR/CD3 and CD68 revealed by FACS in fresh breast cancer samples.

### Identification and Confirmation of TCR+ Macrophages in TNBC tumors

Interestingly, we found a subset of macrophages specifically expressed T cell marker CD3 (Fig. 5c and Supplementary Fig. 3a), as well as the constant region of TCR α/β chains (Fig. 5c), which suggested that these are macrophages bearing α/β TCR (or TCR^+^ macrophages, the same hereinafter). To further confirm that these macrophages expressed paired productive TCRs that can exert antigen recognition function, we reconstructed the TCRs in these macrophages through the scRNA-seq data. Totally, we identified 382 TCR^+^ macrophages with paired productive α and β chains in 13 of the 14 patients, making up 14.28% of the total amount of macrophages, with the proportion ranged from 0.25% to 33.52% across patients (Supplementary Fig. 3b). The percentages of TCR^+^ macrophages in MΦ1 to MΦ6 cluster ranged from 1.97% to 19.67% (Supplementary Fig. 3c). We considered these cells as macrophages rather than T cell for they clustered with other macrophages. In these cells, the expression levels of CD3 and TCR genes were between those for T cells and macrophages, while the macrophage markers had a similar expression level as in macrophages (Fig. 5d). In addition, through reference component analysis, we compared the transcriptome signature of TCR^+^ macrophages with that of the reference bulk transcriptomes summarized by Li *et al*. (44). We found that these cells were more similar to monocytes and macrophages, and were significantly different from T cells, B cells and NK cells (Supplementary Fig. 3e).

Next, we sought to validate the existence of TCR^+^ macrophages. To verify the accuracy of TCR assembly and the expression of intact productive TCR α/β chains, we conducted Sanger sequencing validation in 10 selected single cells (including 5 macrophages and 5 T cells). The whole V region and 50 bp of the 5’ C region of α and β chains were successfully amplified in 9 and 7 cells, respectively by 5’race PCR in corresponding cDNA libraries. The Sanger results of all these amplicons were compared to the assembled TCR α/β chains of the corresponding cells as well as the VJ references from IMGT database. In V/J region, all amplified sequences reached more than 99.5% identity with IMGT references. And in the length of the amplicon, all assembled sequences reached more than 97% identity with corresponding sanger sequences (Supplementary Table 9). To verify the co-expression of CD68 and CD3/TCRα, we conducted multicolor immunofluorescence assays on the tumor specimens of our study cohort. We observed the co-localization of CD68 and CD3/TCRα on the cell membrane in tumor infiltrated immune cells (Fig. 5e and Supplementary Fig. 3d), suggesting the presence of TCR^+^ macrophages. Additionally, we extended our validation by performing FACS analysis using tumor infiltrated immune cells isolated from fresh breast tumors of additional patients independent of our cohort. We found 26.58% and 25.96% of the tumor infiltrated CD68^+^ cells co-expressed CD3 and TCR, respectively (Fig. 5f). Taken together, these results reconfirmed that there is a subset of TCR^+^ macrophages in TNBC microenvironment.

### Potential Function of TCR+ Macropahges Revealed by Uniquely Expressed Genes and TCR-based Analysis

To reveal the potential function of TCR^+^ macrophages, we investigated the differentially expressed genes compared with TCR^−^ macrophages. Notably, significantly up-regulated genes included *LCK*, *LAT*, *FYN*, and *ICOS*, which were necessary for TCR signaling in T lymphocytes, and *GZMA*, and *GZMB*, which were involved in the cytotoxic effect (Fig. 6a, b and Supplementary Table 10). Gene set enrichment analysis showed that the up-regulated genes in the TCR^+^ macrophages (log_2_FC ≥1) were enriched in TCR signaling pathway, MAPK pathway, JAK-STAT signaling pathway, and nature killer cell mediate cytotoxicity pathway (Fig. 6c). These findings illuminated that TCR signaling might actually be activated in these TCR^+^ macrophages. To verify the activation of TCR signaling in TCR^+^ macrophages, we examined the phosphorylation state of ZAP-70, a main switch controlling the activation of TCR signaling pathways (45). Phosphorylated ZAP-70 in TCR^+^ macrophages were observed in tumor specimens from multiple TNBC patients (Fig. 6d). These results demonstrated that a fraction of macrophages in TNBC could not only express TCRs but also might transduce TCR downstream signals, and thus have the potential to activate T cell function associated gene expression.

**Figure 6.**
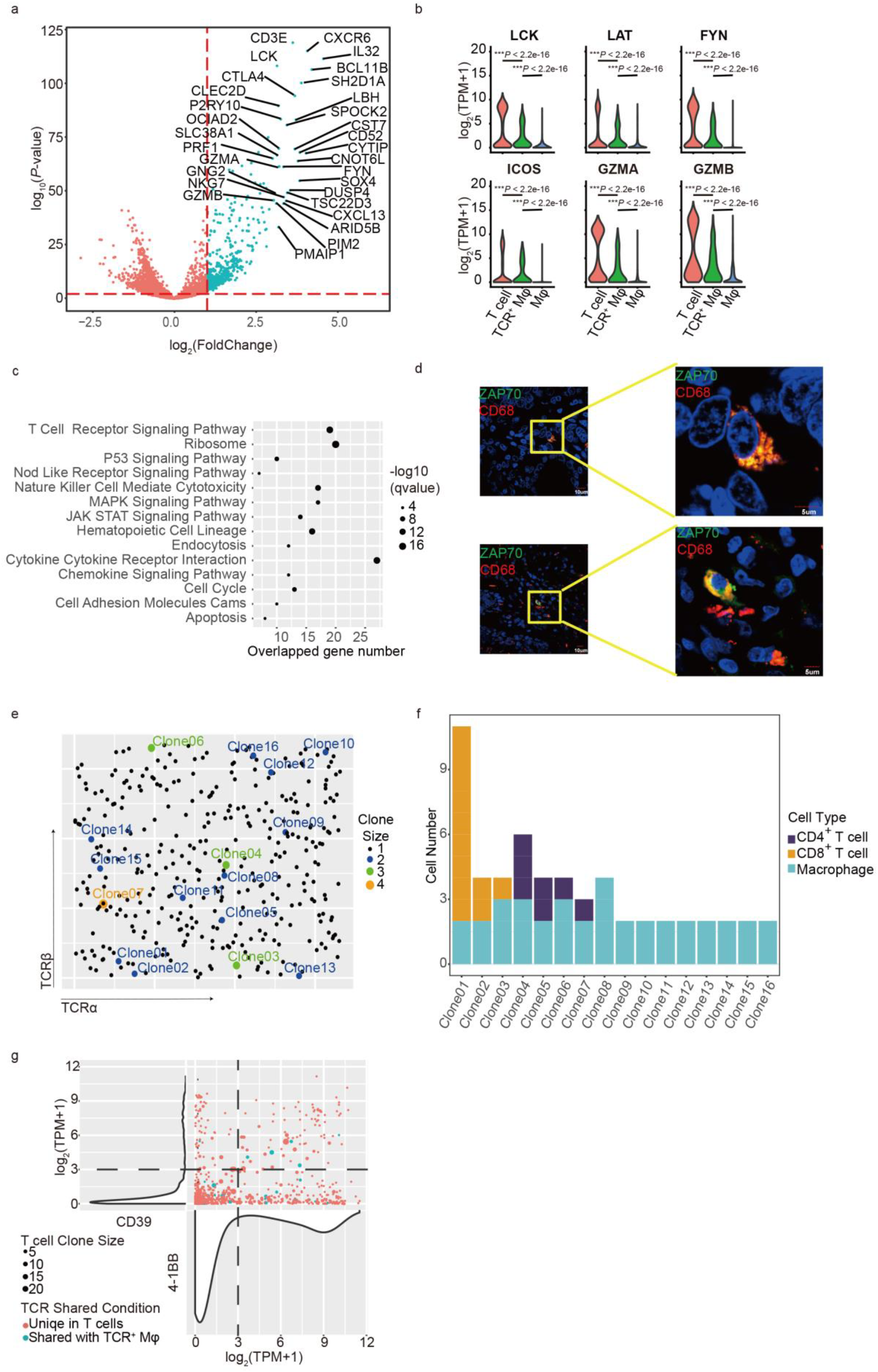
Activation of TCR signaling pathways and TCR diversity in TCR^+^ macrophages. **a**, Volcano plot shows differentially expressed genes in tumor-infiltrating TCR^+^ macrophages compared with TCR^−^ macrophages. Each blue dot represents a gene with *P*-value < 0.01 and log_2_FC ≥ 1. Genes with *P*-value < 0.01 and log_2_FC ≥ 3 are marked with gene names. **b**, The expression (log_2_(TPM+1)) of *LCK*, *LAT*, *FYN*, *ICOS, GZMA*, and *GZMB* are significantly up-regulated in the TCR^+^ macrophages comparing to TCR^−^ macrophages. **c**, Gene set enrichment analysis in KEGG pathway for DEGs with *P*-value < 0.01 and log_2_FC ≥ 1. Selected pathways with *Q*-value < 0.01 are shown in the bubble plot. **d**, Immunofluorescence shows the co-expression of phosphorylated ZAP-70 and CD68 in paraffin sections. **e**, TCR diversity of TCR^+^ macrophages. The x-axis ranked by TCR α chains while the y-axis ranked by β chains. Each point represents a TCR clonotype (TCR αβ pair), colored according to TCR^+^ macrophages’ clone size. Clones with more than one cell are denoted with a given clone number. **f**, The expanded TCR^+^ macrophage clones in **e** share identical TCR with CD4^+^/CD8^+^ T cells. **g**, Average CD39 and 4-1BB expression (log_2_(TPM+1)) of CD8^+^ T cell clones. Each dot represents a CD8^+^ T cell clone. Each dot’s radius depends on the CD8^+^ T cell clone size. CD8^+^ T cell clones sharing TCRs with macrophages are colored by blue. The expressions of CD39 and 4-1BB among CD8^+^ T cells are showed by violin plots on left and below panel, respectively. The high and low expression group of CD39 and 4-1BB are divided by log_2_(TPM+1) = 3.

Moreover, we investigated the characteristics of TCR repertoire of TCR^+^ macrophage. Similar to T cells, macrophages’ TCR repertoire displayed high diversity and individual specificity. Totally, we identified 361 TCR clonotypes. Interestingly, we observed 16 TCR^+^ macrophage clones consisting of at least 2 macrophages, indicating that TCR^+^ macrophages also exhibited clonal expansion (Fig. 6e). Besides, we found that substantial TCR^+^ macrophages shared TCRs with T cells. In total, we identified 61 TCR clonotypes that were shared between TCR^+^ macrophages and CD4^+^/CD8^+^ T cells (Supplementary Fig. 4a, b), of which 7 also existed in macrophage clones (Fig. 6f). Interestingly, we identified 11 public TCR pairs not only shared among macrophages and T cells, but also shared among patients TNBC010 and TNBC012, suggesting that they might recognize common antigens.

We were curious about the antigens these TCR^+^ macrophages target. Recent studies have suggested the existence of bystander CD8^+^ TILs in tumor microenvironment, whose TCRs could only recognize epitopes unrelated to cancer. To investigate whether the TCRs expressed by macrophages also include TCRs that recognize tumor antigen and TCRs that recognize epitopes unrelated to cancer, we looked into the TCRs shared by CD8^+^ TILs and macrophages. We classified the CD8^+^ clones into 4 groups according to the mean expression of *4-1BB* and *CD39* of the clones: 4-1BB^high^CD39^high^, 4-1BB^high^CD39^low^, 4-1BB^low^CD39^high^, and 4-1BB^low^CD39^low^. As the bystander CD8^+^ TILs lacked *CD39* expression (46), and the activated T cells were often marked by high expression of *4-1BB*, we speculated that the 4-1BB^high^CD39^high^ CD8^+^ T cells might recognize and be activated by tumor antigens, and the 4-1BB^low^CD39^low^ T cells might not. We found that in both of the above two groups, there were T cells that shared TCRs with macrophages, indicating that TCRs expressed by macrophages might include ones that might recognize tumor antigens (shared with the 4-1BB^high^CD39^high^ group) and ones that might not (shared with the 4-1BB^low^CD39^low^ group) (Fig. 6g).

## DISCUSSION

Despite major advances in cancer immunotherapy, our ability to understand mechanisms of action or predict efficacy is confounded by the heterogeneous composition of immune cells within tumors. The deep transcriptome data including the complete TCR information for 9,683 individual immune cells provided a comprehensive understanding and multi-dimensional characterization of TNBC infiltrated immune cells. The high quantity and quality of single-cell data allowed us to identify 22 immune cell clusters, including newly reported immune cell subsets, such as CD8^+^CXCL8^+^ naïve T cells and TCR^+^ macrophages. Full-length TCR based analysis enabled us to depict cell clonal expansion patterns and cell lineages. In addition, the sequencing depth in our study assured reliable profiling of cytokines and transcription factors in each cell, allowing a detailed characterization of various immune cell subsets.

Single-cell RNA sequencing or mass cytometry studies of breast tumors and other solid tumors have demonstrated the diversity of tumor-associated immune cell populations, which is supported by our observations in TNBC (17, 47). In this study, we focused our interest on TNBC because it is a breast cancer subtype associated with rapid progression and poor survival once metastasized, although initial responds to chemotherapy and targeted agents sometimes happen. Immune checkpoint blockade therapy has shed light on TNBC treatment by slowing down disease progression and prolonging survival in selected patients (21). A comprehensive knowledge of the TNBC immune ecosystem is pivotal for designing strategies to improve response to immunotherapy in this disease.

The level of tumor-infiltrated T cells and their characteristics are reported to be associated with outcomes of several cancers (9, 10, 48). In our study, we identified T cells with different functional states and gene expression profiles. Psedotime analysis demonstrated the T cell functional state transition from activation to exhaustion along the developmental trajectory. We found the cluster T5, possibly representing cells in a transitional state from effector to exhaustion, indicating a “pre-exhaustion” state of T cells. The “pre-exhaustion” state of T cells have also been suggested in previous studies of NSCLC and HCC tumors (30, 31). Compared to effector T cells, “pre-exhaustion” T cells were characterized by lower expression of cytotoxic markers and elevated expression of exhaustion markers. Further, we found the “pre-exhaustion” signature correlated with better survival in a patient cohort when compared with the signature of exhausted T cell clusters. Thus, identifying these “pre-exhaustion” state T cells and finding ways to reverse these cells to effector-like or preventing them progression into exhaustion could be a potential strategy of immunotherapy.

We reported here a novel T cell subset of CD8^+^CXCL8^+^ naïve T cells (T2), whose signature was linked to unfavorable prognosis. CXCL8 producing CD4^+^ naïve T cells exist in human peripheral blood, mainly in infants (36). Crespo et al. found CXCL8 producing naive CD4^+^ T cells could mediate neutrophil migration in vitro and promotes primary ovarian cancer growth in animal model (37). Consistently, we found the signature of CD8^+^CXCL8^+^ naïve T cell cluster predictive of poor survival in a METABRIC patient cohort. Further DEG and pathway enrichment analysis indicated that tumors showing high T2 signature might progress through mediating leukocytes migration and activating the MAPK/ERK pathways. Taken together, the enrichment of CXCL8^+^CD8^+^ naïve T cells in the TME might contribute to the rapid progression and metastasis of TNBC.

In macrophages, we showed a highly positive correlation between the M1 and M2 signature gene expressions at single cell level. These findings, raising challenges to the customary model of macrophage polarization that M1 and M2 activation states were regarded as mutually exclusive, supported the result of a recent single cell atlas study on breast cancer microenvironment (17). The co-existence and positive correlation of M1/M2 states highlighted the complexity of macrophage functionalities, helping to understand the role of TAM in modulating tumor microenvironment and design macrophage targeted therapies.

Notably, we found a very interesting population of CD3 and TCR positive macrophages in this study. Although TCR^+^ macrophages have been proposed by few previous studies in tuberculous granulomas, lesions of atherosclerosis and carcinomas (49–51), we for the first time observed this macrophage subset in breast cancer microenvironment, and investigated it at single-cell level. We found that TCR^+^ macrophages might widely exist in TNBC tumors, which accounted for averagely 14.28% of the macrophage population in our study, illuminating that this long-neglected macrophage group might be an important component in tumor microenvironment. Single cell RNA-seq data also enabled us to depict the variety of their TCR repertoire. We showed that these TCR^+^ macrophages demonstrated clonal expansion. A fraction of TCRs expressed by macrophages were shared with different subsets of T cells, including Th, Treg and CTLs. Thus, we assume that these macrophages’ TCRs might be shaped by the environment stimuli through a similar manner as the tumor infiltrated T cells, and might target a set of common antigens like T cells. Also, we suggested the activation of TCR signaling pathway in tumor infiltrated TCR^+^ macrophages. These observations raise an attractive perspective that TCR^+^ macrophages in tumor microenvironment may exert part of T cell functions. Whether TCR^+^ macrophages work as a friend or foe with tumor cells in cancer progression deserves in-depth exploration.

In summary, our study highlights the diverse phenotypes of immune cell populations in TNBC microenvironment. Single-cell sequencing data along with TCR information provide us a powerful tool to gain deep insights into the clustering, dynamic, developmental trajectory, and unique signatures of immune cell populations, enabling the discovery of novel immune cell subsets and potential molecular targets for immunotherapy. This immune cell atlas should be a valuable resource for future endeavor to identify clinically relevant cell signatures and predictive markers for precision immunotherapy.

## METHODS

### Isolation of immune cells in human mammary carcinoma

Biopsies with a volume of 0.03-0.3 cm^3^ were mechanically disaggregated to 2 mm^3^ fragments followed by gentle 2 hours enzymatic dissociation with collagenase III (200U/ml), hyaluronidase (100U/ml), Dnase I (0.1mg/ml) in HBSS at 37°C. Following digestion, the cells were washed three times in HBSS. Immune cells were purified by FACS isolation of CD45 positive cells using APC Mouse Anti-Human CD45 (BD Pharmingen, clone: HI30 (RUO), American) as described by the manufacturer.

### Single-cell suspension preparation and RNA extraction

Single-cell suspension preparation was carried out by the protocol previous described (52). RNAs from the single-cells were extracted by a RNeasy plus mini kit (Qiagen) according to the manufacturer’s instructions.

### Single-cell library construction and sequencing

cDNA synthesis and amplification was performed as previous described (52). Amplified cDNA products were purified by 1 × Agencourt AMPure XP beads (Beckman Coulter). A total of 2 ng purified cDNA products from each single-cell were used as the starting amount for library preparation. For the MIRALCS method, amplified cDNA was extracted by an automatic extractor from the chip to 96-well plate and diluted from 200 nl to 5 μl. And 3 μl cDNA products without purification were directly used for library construction. The libraries were prepared by TruePrep™ Mini DNA Sample Prep Kit (Vazyme Biotech) according to the instruction manual and each sample was labelled with a barcode. All of the single-cells were sequenced on BGISEQ500 sequencing platform in a single-end mode.

### Single-cell RNA-seq data processing

The single cell RNA-seq data was produced by BGISEQ-500 in SE100 single-end mode. First, we used Cutadapt (53) to filter reads with adaptors, reads with more than five Ns and reads with low quality. Second, we used Bowtie setting parameters “-q --phred33 --sensitive --dpad 0 --gbar 99999999 --mp 1,1 --np 1 --score-min L,0, −0.1” to map the cleaned reads to the Ensembl hg38 human transcriptome. Then, we used RSEM (RNA-Seq by Expectation Maximization) algorithm to count the number of uniquely mapped reads pair for each gene and then calculate the TPM value for each gene. A readscount matrix and a TPM matrix was made for downstream analysis. We defined genes with TPM > 1 as detected gene and the detected gene number is used for quality control. For each single cell, we calculated the average expression level (log_2_(TPM+1)) of 96 curated housekeeping genes (Supplementary Table 4). To filter out cells with low quality, we set following standard to define high quality cells: 1).Mapping rates ≥ 50%; 2). Fraction of ERCC reads in mapped reads < 50%; 3).Mapped reads ≥ 2M (excluded ERCC reads); 4).Detected gene number ≥ 2500; 5).Housekeeping genes expression level (average log_2_(TPM+1) ≥ 4; 6).Proportions of mitochondrial gene expression value (TPM value) < 10%. Cells defined as high quality were used for downstream analysis. 9683 cells passed the filtering and were used for downstream analysis.

### Unsupervised clustering

We first normalized the TPM value as log_2_(TPM+1). To remove the potential variation caused by individual differences, we further centered the log normalized TPM matrix patient by patient. If no explicitly stated, “normalized TPM” in this study refers to the log normalized and centered TPM value. We used Seurat2.0 (54) to perform unsupervised clustering on the single cells using the normalized TPM matrix as input. We kept genes annotated as “protein coding” for downstream analysis. Next, we considered genes with top 3000 highest standard deviation as highly variable genes excluded genes with average TPM < 1 (not normalized) across all cells to avoid unexpected noise. To regress out the variances caused by cycling cells from our data, we performed“Cell-Cycle Scoring and Regression”recommended by the tutorial of Seurat following the reported protocol (55). In this step, single cells were scored basing on the scoring strategy described by Tirosh et al (32) using a canonical gene list (Supplementary Table 5) and the relationship between gene expression and the cell cycle score was modeled to create a corrected expression matrix used for downstream dimensional reduction. After regressing out the cell cycle affect, we performed PCA on the corrected expression matrix using highly variable genes described above. Following the results of PCA, we selected the first 20 PCs for downstream analysis. Finally, we used the Seurat function “FindClusters” with the “resolution” parameter set to 1.5 to cluster the single cells. As a result, we obtained 22 cell clusters and visualized them by t-SNE.

### Cell type annotation

To annotate our clusters to specific immune cell types (T cells, B cells, Macrophages, DCs, NKs), we selected some classic markers of these main immune cell types (Supplementary Table 6) and plotted their expression level on the t-SNE graph. Moreover, we analyzed differentially expressed genes (DEGs) among clusters to obtain the cluster specific genes of each cluster by R package limma (56) using voom approach. The classic markers with adjusted p value (Benjamini-Hochberg multiple testing correction) < 0.01, log2FC ≥ 3 and expressed by > 50% of the cells (TPM > 1) in that cluster were used to define cell type (Supplementary Table 6). Furthermore, we compared cluster mean expression (log_2_(TPM+1)) to sorted bulk transcriptome profile using RCA (44). The results of marker annotating, DEG and RCA results were considered together to determine the cell type of our clusters. To further validate the clustering and cell type annotation results and regress out doublets and potential noise, we performed RCA on each single cell to annotate each single cell to a specific immune cell type and filtered out cells which were assigned to a different cell type with their cluster. We next identified CD4^+^ and CD8^+^ cells based on the expression of *CD8A*, *CD8B* and *CD4* in the T cell population. Only cells with the average TPM of *CD3D*, *CD3E* and *CD3G* larger than 10 were kept for this analysis. Based on the average TPM of *CD8A* and *CD8B*, one cell was considered as CD8 positive or negative if the value was larger than 7 or less than 1, respectively. Based on the TPM of *CD4*, one cell was considered as CD4 positive or negative if the value was larger than 7 or less than 1, respectively. As a result, cells were classified as CD4^+^CD8^+^ (double positive, DP), CD4^−^CD8^−^ (double negative, DN), CD4^−^CD8^+^ (CD8^+^T), CD4^+^CD8^−^ (CD4^+^T) and other cells that cannot be clearly identified. Furthermore, T1, T2, T3, T4, T5, T6 were defined as CD8^+^ T cluster while T7, T8, T9 as CD4^+^ T cluster according to the cell content of these T cell clusters.

### Cluster marker analysis

In order to define the subtype of T cells and Macrophages, we analyzed the DEGs among T cell clusters and Macrophage clusters separately by the following step: 1). DEGs among clusters were analyzed by R package limma using voom approach. Genes with Benjamini–Hochberg-adjusted F-test *P*-value < 0.01 and log_2_FC ≥ 1 were kept. 2). The area under the curve value (AUC) was calculated for each gene to measure its ability to discriminate one cluster from the remaining clusters: each gene’s log normalized TPM value was treated as a predictor, and cells inside and outside of the cluster were treated as positive and negative instances, respectively. Genes with AUC ≥ 0.5 were kept. 3). The identified DEGs were then categorized to the cluster showing the highest FC and with at least 50% of cells expressing this gene (TPM>0, Table S4).

### DEG Analysis for Exhausted and Non-exhausted CD8+ T Cells

We defined cells in T1 as exhausted CD8+ T cells because there are several classic exhausted T cell markers such as HAVCR2, IDO1 and LAG3 significantly express in these clusters (adjusted p value < 0.01, fold change >= 4). We defined cells in other CD8+ Tcell clusters as non-exhausted T cells. DEGs analysis between exhausted and non-exhausted CD8+ T cells was carried out using the voom method by limma described above with thresholds for adjusted *P*-value <0.01 and fold change >=4.

### DEG Analysis for Treg and Other CD4+ T cells

Cells of T7 and T9 were defined as Treg while T8 as Tconv. We performed DEGs analysis using limma by voom approach described above between Treg and Tconv to get the Treg-specific genes with different threshold (adjusted *P*-value < 0.01 and fold change >= 2).

### Cell developmental trajectory

We used the Monocle2 (29) to order cells in CD8^+^/CD4^+^ T cell clusters in pseudo time. We first used the “relative2abs” function in Monocle to convert TPM into normalized mRNA counts and created an object with parameter “expressionFamily = negbinomial.size” following the Monocle2 tutorial. We used “differentialGeneTest” function to derive DEG from clusters, keeping the genes with *Q*-value <0.01. After all, genes with mean expression ≥ 0.1 (normalized mRNA read counts by “relative2abs” function) in the kept DEGs were used to order the cells in pseudo time.

### DEG analysis for exhausted and non-exhausted CD8^+^ T cells

We defined cells in T1 and T6 as exhausted CD8^+^ T cells and cells in other CD8^+^ T cell clusters as non-exhausted CD8^+^ T cells. DEGs analysis between exhausted and non-exhausted CD8^+^ T cells was carried out using the voom method by limma described above with thresholds for adjusted *P*-value <0.01 and log2FC ≥ 2 (Table S5).

### Survival analysis

The METABRIC (57) dataset were used to evaluate the prognostic effect of signature gene sets derived from cluster. The gene expression and survival data of the METABRIC dataset were accessed using CBioPortal (58). For each signature gene set, we calculated a “gene signature score” for each patient using fold-change values for each gene in the signature (Supplementary Table 1) to weight each gene in calculating the average. Then, patients were grouped into high and low expression groups by the median value of the “gene signature score”. To correct clinical covariates including age and histological grade, we performed Multivariable analyses using Cox proportional hazards survival models and get the Hazard Ratio (HR) and adjusted *P*-value.

### DEGs between patients grouped by the gene signature score of T2

As described in the “Survival analysis”, patients were grouped into high and low expression groups by the median value of the “gene signature score” of T2. DEGs analysis between T2^high^ (n=952) and T2^low^ (n=952) patients was carried out following the best practice for microarray data using limma.

### TCR Analysis

We used TraCeR (59) method to assemble the TCR sequences for each single cell. Tracer can identify the rearranged TCR chains and use kallisto to calculate their TPM values. For every single cell, we rearranged its productive TCR chains by their TPM values. For example, if two TCRα chain were assembled in one single cell and they were both productive, the chain with higher TPM will be defined as TCRα1 while the chain with lower TPM as TCRα2.Non-productive TCR chains were excluded. The same rearrangement was deployed on TCRβ. We kept cells with at least one pair of productive TCRα and TCRβ chain for subsequent analysis. We used a strict standard to define TCR clones: cells with the same TCRα1 and TCRβ1 were considered to be one TCR clone while the expanded clones were defined as clones with at least two cells sharing the same TCRα1 and TCRβ1 in a given cell population.

When analyzing the sharing TCR within CD8^+^ T cells / CD4^+^ T cells or between CD8^+^/CD4^+^ T cells and Macrophages, only cells defined as CD4^−^CD8^+^ T cells or CD4^+^CD8^−^ T cells according to CD4/CD8 expression (see the cell type annotation section) were included in the analysis

### TCR sequence sanger validation

Single cells’ mRNA was reverse transcripted into cDNA during the RNA-seq already. The cDNA was amplificated with TCR primer by three step nest-PCR. First PCR needs 2 × KAPA Ready mix 12.5μM, 10uM IS PCR primer 0.5μM, 10uM CA-out (5’-GCAGACAGACTTGTCACTGG-3’) 1μM, 10uM CB-out (5’-TGGTCGGGGAAGAAGCCTGTG-3’) 1μM, RT mix 10μM, and the amplification steps were 98 °C 3min; 98 °C 15 s, 55 °C 20 s, 72 °C 2 min × 25 cycles; 72 °C 5 min, 4 °C hold. The second PCR has 2 × KAPA Ready mix 12.5μM, 10uM IS PCR primer 0.5μM, 10uM CA-in-N (5’-AGTCTCTCAGCTGGTACACG-3’) 1μM, 10uM CB-in-N (5’-GCAGACAGACTTGTCACTGG-3’) 1μM, the first PCR RT mix 10μM, and the amplification steps were 98 °C 3min; 98 °C 15 s, 60 °C 20 s, 72 °C 2 min × 25 cycles; 72 °C 5 min, 4 °C hold. The third PCR contains 2 × KAPA Ready mix 12.5μM, 10uM IS PCR primer 0.5μM, 10uM AC-rev-1 (5’-GGTACACGGCAGGGTCAGGGTTC-3’) 1μM, 10uM BC-rev-1 (5’-TTCTGATGGCTCAAACACAGCGA-3’) 1μM, the second PCR RT mix 10μM, and the amplification steps were 98 °C 3min; 98 °C 15 s, 60 °C 20 s, 72 °C 2 min × 35 cycles; 72 °C 5 min, 4 °C hold. Then 1% agarose gel electrophoresis was used to recover the target DNA. Connected DNA fragment to pMDTM18-T vector (TaKaRa 6011) at 4 °C overnight. Transformed the product into Trans5α (Transgen, CD201), and plated the bacterial on ager medium with Ampicillin, X-Gal and ITPG at 37°C for 16 h. Picked one white bacterial colony to 20μl ddH2O, then took 1μl to PCR with R primer (5’-CGCCAGGGTTTTCCCAGTCACGAC-3’) 1μl, and F primer 1μl, dNTPs 4μl, r-Taq 1μl, 10× Buffer 5μl, ddH2O 37μl. The amplification steps were 98 °C 5min; 95 °C 30 s, 60 °C 30 s, 72 °C 40 s × 35 cycles; 72 °C 10 min, 4 °C hold. Performed 1% agarose gel electrophoresis to recover the target DNA. The recovered DNA was sequenced by sanger sequencing.

### TCR sanger sequencing result analysis

Sanger sequencing was performed and we obtained the TCR sequence of some of the single cells. We blast the sequence against the sequence assembled by Tracer method and the VDJ sequences in IMGT database to proof the result of Tracer (Supplementary Table 4).

### M1 and M2 signature analysis

M1 and M2 signature genes were provided by Azizi.et al (60) (Table S6). The normalized TPM (see the unsupervised clustering section) was used for analysis. We calculated the mean expression of M1/M2 signature for each cell and calculated the Pearson correlation of M1 and M2 signature.

### Flow Cytometry

5 × 5 × 5 mm tumor samples were resected during the operation and cut into pieces. Added the digestive fluid to the tubes and isolated the tissue into single cells. Dyed he cells with TCR α/β PE (555548, BD), CD68 FITC (333806, Biolegend), CD3 APC (17-0036-42, eBioscience) antibodies, grouped the samples and analyzed with CytoFLEX FCM (Beckman).

### Immunofluorescence

The tissue sections were placed at room temperature for 1h before dewaxing. Soaked the sections in dimethylbenzene for 10min and change the dimethylbenzene for 10min more. Immersed the sections in 100% −95%-85%-75% ethanol for 5min each, washed by PBS for 5min twice. Bathed the sections in 0.01M sodium citrate buffer solution (pH 6.0) at 95 °C for 15min, washed with PBS for 5min. Added 5% goat serum to the slides and sealed at room temperature for 30 minutes. Added CD3 (ZSGB-BIO, ZA-0503), CD68(ZSGB-BIO, ZM-0464), ZAP70(Abcam, ab76306), TCR α (Abcam, ab18861), primary antibody (1:200) overnight at 4°C, washed with PBS for 5min three times. The fluorescently labeled secondary antibody (1:100) Dylight 488(Abbkine, A23220) and Dylight 549(Abbkine, A23310) was incubated for 1 h at room temperature in the dark. Bathed in PBS for 5min twice, dropped DAPI (Abcam, ab104139) and applied cover slip carefully. Sealed with nail polish and dried at room temperature. The immunofluorescence pictures were taken by laser scanning confocal microscope (Olympus, FV1000) with 405nm, 488nm, 559nm and 633nm.

## Availability of data and materials

The Single-cell RNA sequencing data in this study are available in the CNGB Nucleotide Sequence Archive (CNSA: https://db.cngb.org; accession number: CNP0000286).

## Acknowledgements

We thanked Fei Wang for his technical support in single cell TCR sequencing and validation. We thanked Dr. Song Gao, Dr. Leo J Lee and Dr. Weimin Zhang for their valuable advices for this paper. This project was supported by grants from Joint Fund of the National Natural Science Foundation of China and Natural Science Foundation of Guangdong Province (U1601224), National Natural Science Foundation (81602342, 81602313), Natural Science Foundation of Guangdong Province (2017A030313466) and Shenzhen Municipal Government of China (No. 20170731162715261)

## Author contributions

S. W., B.L., Q.Z., S.Q. and R.H. conceived the project. Y.L., L.Z. and X.L. designed the experiments. Y.L., L.Z., X.L., K.Z., M.H., J.L. and F.W. performed the experiments. Q.L., W.X., J.T., X.X., W.W., Y.S. and Z.Y. collected clinical samples. S.Q., Z.Z., S.L analyzed the sequencing data. S.W., B.L., S.Q., R.H., Z.Z., Y.L., Z.L. wrote the manuscript with all authors contributing to writing and providing feedback.

